# Comprehensive Analysis of Neutrophil Immunomodulatory Properties and FcR Dynamics in Health and HIV-1 Infection and Therapy

**DOI:** 10.1101/2024.02.13.580065

**Authors:** Soledad Marsile-Medun, Manon Souchard, Daouda Abba Moussa, Valérie Lorin, Hugo Mouquet, Elisa Reynaud, Rayane Dibsy, Edouard Tuaillon, Delphine Muriaux, Giang Ngo, Martine Pugnière, Mar Naranjo-Gomez, Mireia Pelegrin

**Author notes:** **Corresponding authors:** Mar Naranjo-Gomez and Mireia Pelegrin, Institute of Regenerative Medicine and Biotherapy of Montpellier, UMR1183, 80, Avenue Agustin Fliche, 34293 Montpellier Cedex 5, France. equally contributed (Mar Naranjo-Gomez, Mireia Pelegrin).

## Abstract

Neutrophils are innate immune cells with key immunomodulatory functions. In a murine retroviral model, we previously showed their essential role in promoting protective immunity during antiviral antibody therapy via Fc–Fcγ receptor (FcγR) interactions. Here, we investigated the immunomodulatory properties of neutrophils in the context of HIV-1 infection and therapy through a comprehensive analysis of their functional activation and the regulation of FcγR expression. Neutrophils from healthy donors (HD) and people living with HIV-1 (PLWH) were stimulated with TLR ligands, free HIV-1, immune complexes (ICs) formed with broadly neutralizing antibodies (bNAbs), or pro-inflammatory cytokines (TNFα, IFNγ). In response, they secreted various cytokines and chemokines that can recruit and activate immune cells in a stimulus-dependent manner. Compared to TLR agonist and cytokine activation, HD neutrophils showed limited cytokine production in response to free HIV-1 or ICs alone as well as a minimal FcγR modulation. However, PLWH neutrophils showed heightened responsiveness to microbial stimuli linked to HIV-1 pathogenesis, secreting higher levels of IFNγ, CXCL1, CCL2, CCL3, and CCL4. They also expressed higher levels of two activating FcγRs (FcγRI and FcγRIIIb), as well as CD11b, CD63, CXCR4, and PD-L1, indicating an altered activation state. These findings highlight the influence of the inflammatory milieu on neutrophil function and FcγR regulation in HIV-1 infection and mAb-based therapies.

## Introduction

Neutrophils are the most abundant population of circulating white blood cells and are massively and rapidly recruited following infection. Through diverse mechanisms such as phagocytosis, releasing of reactive-oxygen species (ROS), trapping of the pathogenic agent by neutrophil extracellular traps (NET) (known as NETosis) (1), neutrophils can limit and counteract invading pathogens. Moreover, immunomodulatory properties of neutrophils have been described in recent years, showing their ability to shape the adaptive immune response (2,3). Neutrophils migrate from the site of infection to draining lymph nodes where they interact with multiple immune cells through the production of several cytokines and chemokines (3), making neutrophils key cells in the orchestration of innate and adaptive immunity (3,4). However, the immunomodulatory role of neutrophils in antiviral immunity is still ill-understood, including in the context of persistent viral infections such as HIV-1, as well as in the frame of antiviral monoclonal antibody (mAb)-based immunotherapies. Noteworthy, it has been reported that the functional activation of neutrophils is compromised upon HIV-1 infection which leads to impaired effector functions (i.e. phagocytosis, ROS release, …) and increased apoptosis (5,6). In addition, more recent studies show that neutrophils in people living with HIV-1 (PLWH) are highly activated and may contribute to ongoing chronic inflammation in HIV (7). However, much less is known on the effect of HIV-1 infection on their cytokine and chemokine secretion profile, as well as on the modulation of Fcγ receptors (FcγRs) expressed at their surface, which is a key point in the context of antibody therapy.

The discovery of potent broadly neutralizing antibodies (bNAbs) against HIV-1 has made passive immunotherapy a potential strategy for the prevention and treatment of HIV-1 infection (8). Importantly, recent clinical data have demonstrated the efficacy of several anti-HIV-1 bNAbs to control viremia when administered to PLWH, as well as to induce adaptive immune responses (vaccinal effect) (reviewed in 9). This has given strong support to the idea that bNAbs could broaden the therapeutic arsenal against HIV-1 infection (10–14). Beyond their neutralization capacity through the binding of their Fab fragment to viral antigens, the biological activity of mAbs is also mediated by the Fc moiety upon interaction with the complement system and FcγRs expressed by many cells of the immune system (15,16). This allows to clear free virions from the blood as well as to guide host immune effector cells to kill infected cells by several Fc-mediated mechanisms (i.e. antibody-dependent cellular phagocytosis (ADCP), antibody-dependent cell-mediated cytotoxicity (ADCC), …). Furthermore, mAbs can form immune complexes (ICs) *via* binding with either virions or infected cells, and the subsequent engagement of FcγRs by ICs has immunomodulatory effects leading to the induction of protective immunity. We have previously identified different FcγR-expressing immune cells involved in the vaccinal response during mAb-based therapies (reviewed in 17,18). By using a mouse model of retroviral infection, we have shown a key role for ICs in enhancing antiviral immune responses by multiple innate immune cells such as dendritic cells (DC) (19,20), neutrophils (21,22), monocytes (22) and NK cells (23) in a FcγR-dependent manner. Notably, our findings revealed a key immunomodulatory role for murine neutrophils in the induction of protective immunity upon mAb-therapy (21,22). This occurs through the secretion of multiple cytokines and chemokines upon activation by free virions and ICs. With the aim of extending our observations to the context of HIV-1 infection, here we addressed the immunomodulatory properties of human neutrophils upon activation by different microbial stimuli associated with HIV-1 infection and mAb-based therapy.

Studying the functional properties of neutrophils presents significant challenges due to their short half-life, fragility, and susceptibility to activation during the purification process. Most studies on their immunomodulatory properties focus on a limited set of immune parameters, often relying on gene expression data and lacking a comprehensive assessment of their functional activation and FcγR modulation. Furthermore, existing data are fragmented and difficult to compare, as variations in purification methods can influence neutrophil activation status. Additionally, variations in pathophysiological conditions (e.g., healthy state or infections caused by viruses, bacteria, or fungi) or therapeutic interventions affecting neutrophil donors introduce further variability, complicating the interpretation and comparison of findings. Here, we conducted a comprehensive and systematic investigation of neutrophil activation, analyzing multiple immune receptors and a broad panel of chemokines and cytokines in neutrophils isolated from HD and PLWH upon activation by various stimuli associated with HIV-1 infection. We observed that neutrophils exhibited a limited capacity to secrete cytokines and chemokines in response to *bona fide* HIV-1 (free virus or ICs) compared to virus-mimicking compounds such as TLR agonists, commonly used as viral surrogates. Our findings also reveal distinct phenotypic and functional differences in neutrophils from PLWH compared to those from HD, including heightened responsiveness to microbial/inflammatory stimuli and upregulated expression of activating FcγRs. These alterations may influence the effectiveness to bNAb therapies. By providing a comprehensive analysis of neutrophil activation and FcγR modulation in response to various HIV-1–related stimuli, this study offers valuable insights into neutrophil-mediated mechanisms that could influence the outcomes of antibody-based therapeutic strategies—either alone or in combination with TLR agonists and/or compounds targeting neutrophil immunomodulatory functions.

## Results

### Different cytokine/chemokine secretion profile and FcγR cell surface levels in neutrophils stimulated by TLR agonists or pro-inflammatory cytokines

First, we assessed the effect of TLR agonists and inflammatory cytokines (TNF-α and IFN-γ) on the phenotypic and functional activation of human neutrophils isolated from HD. This aimed to determine whether and how the microenvironment resulting from infection might be determinant for modulating the immune response, both in terms of chemokine/cytokine secretion profile and of FcγR expression pattern, as observed in our mouse studies (22). As for TLR agonists, we mainly focused on TLR4 (lipopolysaccharide (LPS)) and TLR7/8 agonists and Resiquimod (R848), due to their role in immune activation associated with viral sensing (TLR7/8) and to the bacterial translocation observed in PLWH (i.e. LPS binding to TLR4). To assess the phenotypic and functional activation of neutrophils we used the following read-out: (i) the upregulation of the cell surface markers CD11b and CD66b, (ii) the modulation of the expression of the main FcγRs constitutively expressed on neutrophils: FcγRIIa (CD32a) and FcγRIIIb (CD16b), and (iii) the cytokine and chemokine secretion profile (**Figure 1**). Flow cytometry analysis was performed on the viable fraction of CD15^+^/CD11b^+^ cells (**Figure 1A**). While life span of neutrophils has been reported to be short (only few hours), specific culture conditions allow extending it. Upon TLR4 and TLR7/8 agonist stimulation during 16-20h, neutrophil survival was around 65 and 75% in average, respectively (**Figure 1B**). TLR7/8 agonist led to significant upregulation of CD11b as compared to unstimulated neutrophils and TLR4 agonist stimulation, while TLR4 agonist led to significant upregulation of CD66b as compared to unstimulated neutrophils, with no significant differences between TLR agonists (**Figure 1C)**. As for FcγR expression, neutrophil stimulation by either TLR agonist led to decreased expression of the CD32a receptor, as well as a significant decrease in CD16b cell surface levels by TLR4 activation, as compared to unstimulated neutrophils (**Figure 1C)**. We next assessed the functional activation of TLR-stimulated neutrophils. Secretion of soluble cytokines (IL-1β, Il-6, TNF-α, IFN-λ1, IL-8, IL-12p70, IFN-α2, IFN-λ2/3, GM-CSF, IFN-β, IL-10, IFN-γ) and chemokines (CXCL10, CCL11, CCL17, CCL2, CCL5, CCL3, CXCL9, CXCL5, CCL20, CXCL1, CXCL11, CCL4) were assayed from cell-free culture supernatants of *in vitro* stimulated neutrophils using bead-based immunoassays. The assessment of neutrophil secretome revealed that both stimuli triggered the secretion of IL-8 and several chemokines (i.e. CCL3, CCL4, CXCL1, CXCL10 and CCL2) albeit to a different extent (**Figure 1D)**. Due to high donor-dependent variability of chemokine secretion, no statistical differences were observed between TLR4 and TLR7/8 activation (**Figure 1D**). However, TLR4 stimulation led to the secretion of IFN-λ2/3 in 3 out of 7 samples, while TLR7/8 stimulation did not. No detectable release of the other cytokines or chemokines was observed under our experimental conditions.

**Figure 1.**
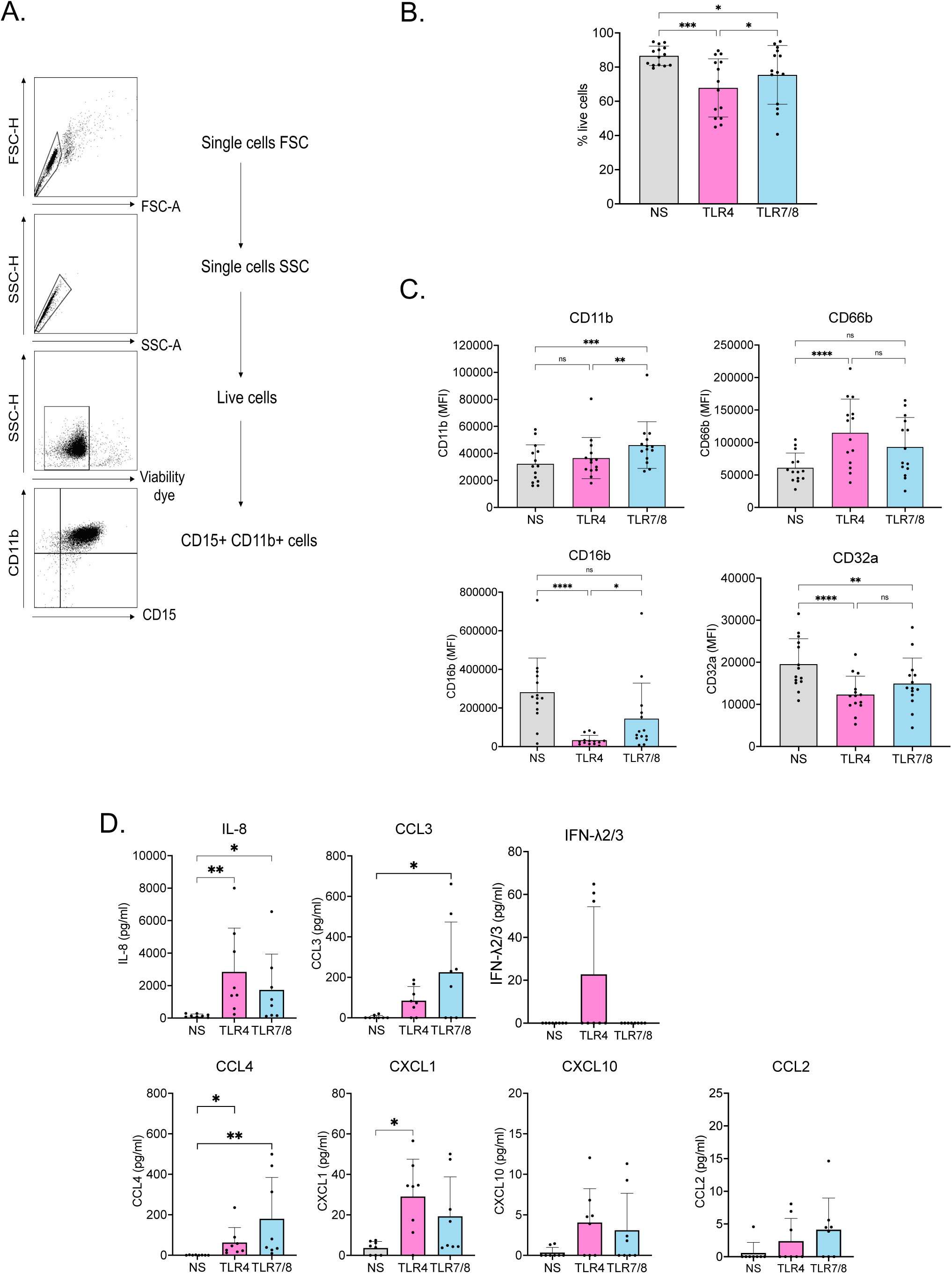
Activation of human neutrophils stimulated *in vitro* by Toll-like receptor (TLR) agonists. Neutrophils from HD were activated overnight by TLR4 (LPS) or TLR7/8 (R848) agonists. Activated neutrophils and corresponding cell culture supernatants were collected to evaluate neutrophil viability and phenotypic activation via flow cytometry, as well as to analyze their secretome using bead-based immunoassays. (A). Gating strategy used for all experiments. (B-C) Viability (B) and phenotypic activation (C) by monitoring cell surface markers using flow cytometry. (D) Cytokines and chemokines secretion profile. Secretion of soluble cytokines (IL-1β, Il-6, TNF-α, CXCL10, IFN-λ1, IL-8, IL-12p70, IFN-α2, IFN-λ2/3, GM-CSF, IFN-β, IL-10, IFN-γ) and chemokines (IL-8, CXCL10, CCL11, CCL17, CCL2, CCL5, CCL3, CXCL9, CXCL5, CCL20, CXCL1, CXCL11, CCL4) was assayed from cell-free cell culture supernatants. Only cytokines/chemokines detected in cell culture supernatants are represented. Values are mean +/− SD of 14 (B-C) and 8 (D) independent measurements. Significance was assigned as follows: ∗p < 0.05, ∗∗p <0,01, ∗∗∗p < 0.001 and ∗∗∗∗p < 0.0001 NS: non-stimulated.

We also assessed the viability, phenotype, and functional activation of human neutrophils stimulated with TNF-α and IFN-γ (**Figure 2**). These cytokines were chosen based on evidence linking virus-driven inflammation, including TNF-α/IFN-γ signatures, to the development of improved antiviral responses in spontaneous HIV controllers (24). At the concentrations used, TNF-α led to a similar neutrophil survival than IFN-γ (75% and 80%, respectively) (**Figure 2A**). TNF-α induced a significant upregulation of CD66b as well as a significant decrease of CD32a and CD16b cell surface levels. In contrast, the expression levels of these activating markers and FcγRs were not significantly altered in neutrophils activated by IFN-γ (**Figure 2B**). Notably, IFN-γ stimulation led to a significant increase in the surface expression of the inducible FcγRI (CD64) (15), an effect not observed with TNF-α (**Supplementary Figure 1)**. Thus, significant differences between IFN-γ and TNF-α activation were observed for most of the cell surface markers evaluated. The assessment of neutrophil’s secretome revealed that TNF-α activation induced the secretion of several cytokines (IL-8, IL-1β, IFN-λ2/3, IFN-λ1 and IFN-γ) and chemokines (CCL3, CCL4 and CXCL1) (**Figure 2C**). The secretion of IL-8, CCL3, CCL4, and CXCL1 was significantly higher compared to unstimulated neutrophils. Increased levels of IL-1β, IFN-λ2/3, IFN-γ, and CCL2 were also observed in approximately half of the samples tested, although these changes did not reach statistical significance (**Figure 2C**). In contrast, IFN-γ activation did not induce detectable cytokine or chemokine release, except for a low level of IL-8 secretion.

**Figure 2.**
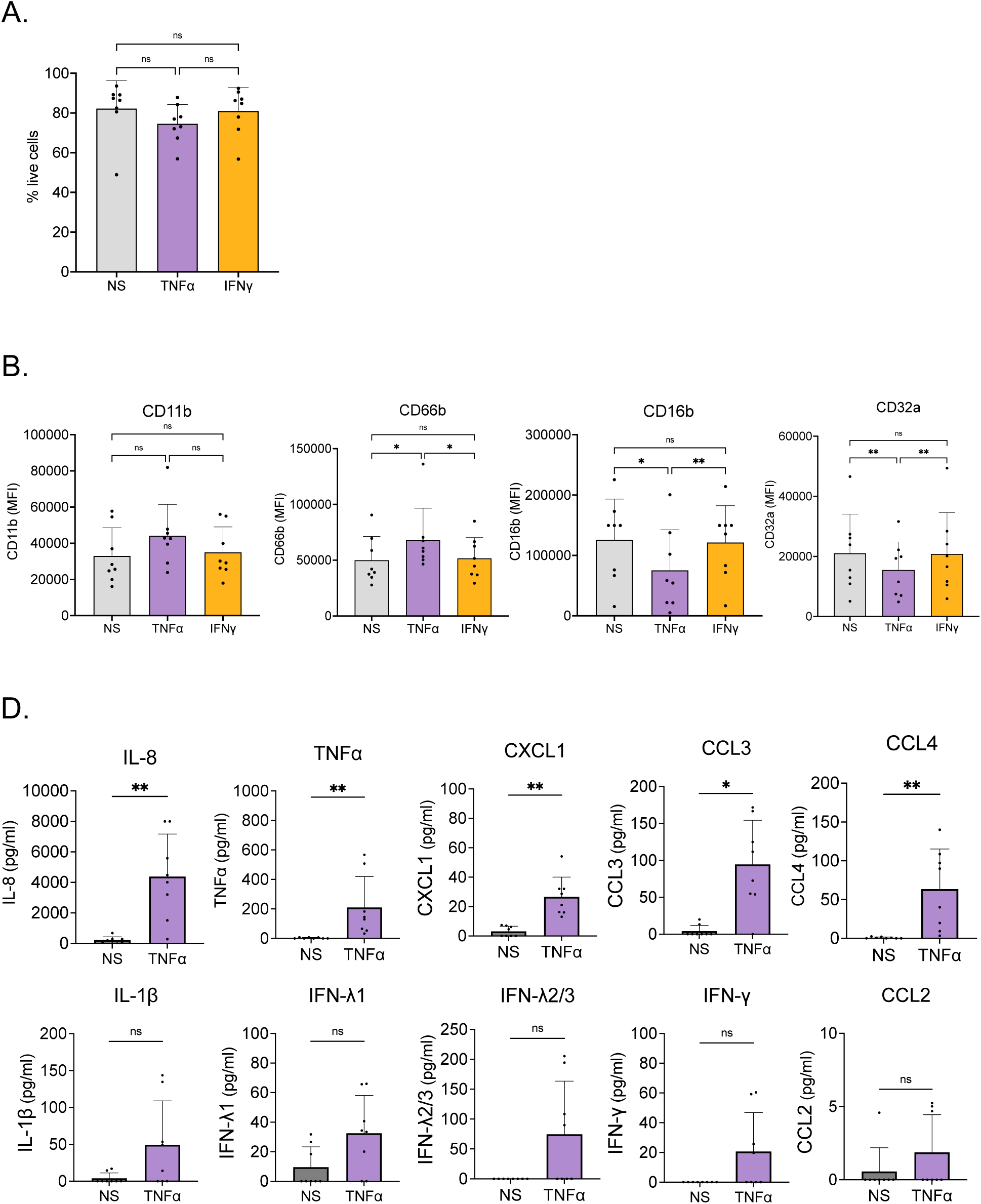
Activation of human neutrophils stimulated *in vitro* by TNF-α or IFN-γ. Neutrophils from HD were activated overnight by TNF-α at 1 ng/mL or IFN-γ at 50 ng/ml. Activated neutrophils and corresponding cell culture supernatants were collected to evaluate neutrophil viability and phenotypic activation via flow cytometry, as well as to analyze their secretome using bead-based immunoassays. (A-B) Viability (A) and phenotypic (B) activation assessed by monitoring cell surface markers. (C) Cytokines and chemokines secretion profile was performed as in Figure 1 and is represented for each cytokine/chemokine detected upon TNF-α. IFN-γ activation did not induce detectable cytokine or chemokine release, except for a low level of IL-8 secretion, and is therefore not shown in panel C. Values are mean +/− SD of 8 independent measurements. Significance was assigned as follows: ∗p < 0.05, and ∗∗p < 0.01. NS: non-stimulated.

Overall, these results highlight a stimulus-specific effect on neutrophil activation, both phenotypically (FcγR and activation marker levels) and functionally (secretome). Furthermore, by studying multiple parameters, these results enabled to establish a comprehensive and comparative landscape of neutrophil activation by these stimuli (some of them used as a surrogate of viral activation) as a basis for comparison with neutrophil activation by HIV-1 and ICs. This is important to highlight as neutrophil activation by TLR agonists and cytokines has been partially addressed in previous studies (25–29). However, the available data are scattered, limited to a narrow set of parameters, and derived from diverse experimental conditions.

### Neutrophil FcγR Expression is maintained upon HIV-1 stimulation

We next examined the modulation of the expression of FcγRs in presence of HIV-1 virions, free or in the form of ICs. This is important to assess, as in the context of HIV-1 infection and antibody therapy, the expression of FcγRs might affect the activation of neutrophils by therapeutic antibodies. ICs were made using the anti-HIV-1 bNAb 10-1074, which is currently being tested in clinical trials and has been shown to induce vaccinal effects (9,30). The 10-1074 bNAb was chosen due to its higher affinity to the FcγRs CD16b and CD32a as compared to another bNAbs also used in the clinic (3BNC117) (**Supplementary Figure 2**). Activation by HIV-1 virions, ICs and free bNAb had no significant effect on neutrophil survival, which was maintained around 85%, similarly to unstimulated neutrophils (**Figure 3A**). The stimulation with HIV-1 virions and ICs led to mild but significant upregulation of CD11b and CD66b as compared to unstimulated neutrophils (**Figure 3B**). Both stimuli induced comparable neutrophil activation — as evidenced by increased expression of activation markers (**Figure 3B**) and cytokine/chemokine secretion (**Figure 3C**). However, no significant downregulation of CD16b expression was observed in cells stimulated with HIV-1 or ICs compared to unstimulated cells. A significant decrease in CD32a expression was observed in IC-activated neutrophils (**Figure 3B**), which may reflect a specific interaction between ICs and this FcγR receptor. Importantly, these findings demonstrate that despite neutrophil activation by HIV-1 and ICs, the expression of FcγRs—particularly CD16b—is largely preserved. This limited modulation of FcγRs suggests that neutrophil effector and immunomodulatory functions mediated through antibody-dependent mechanisms remain intact, highlighting a potentially favorable immunological landscape for Fc-mediated immunity and therapeutic strategies against HIV-1 infection.

**Figure 3.**
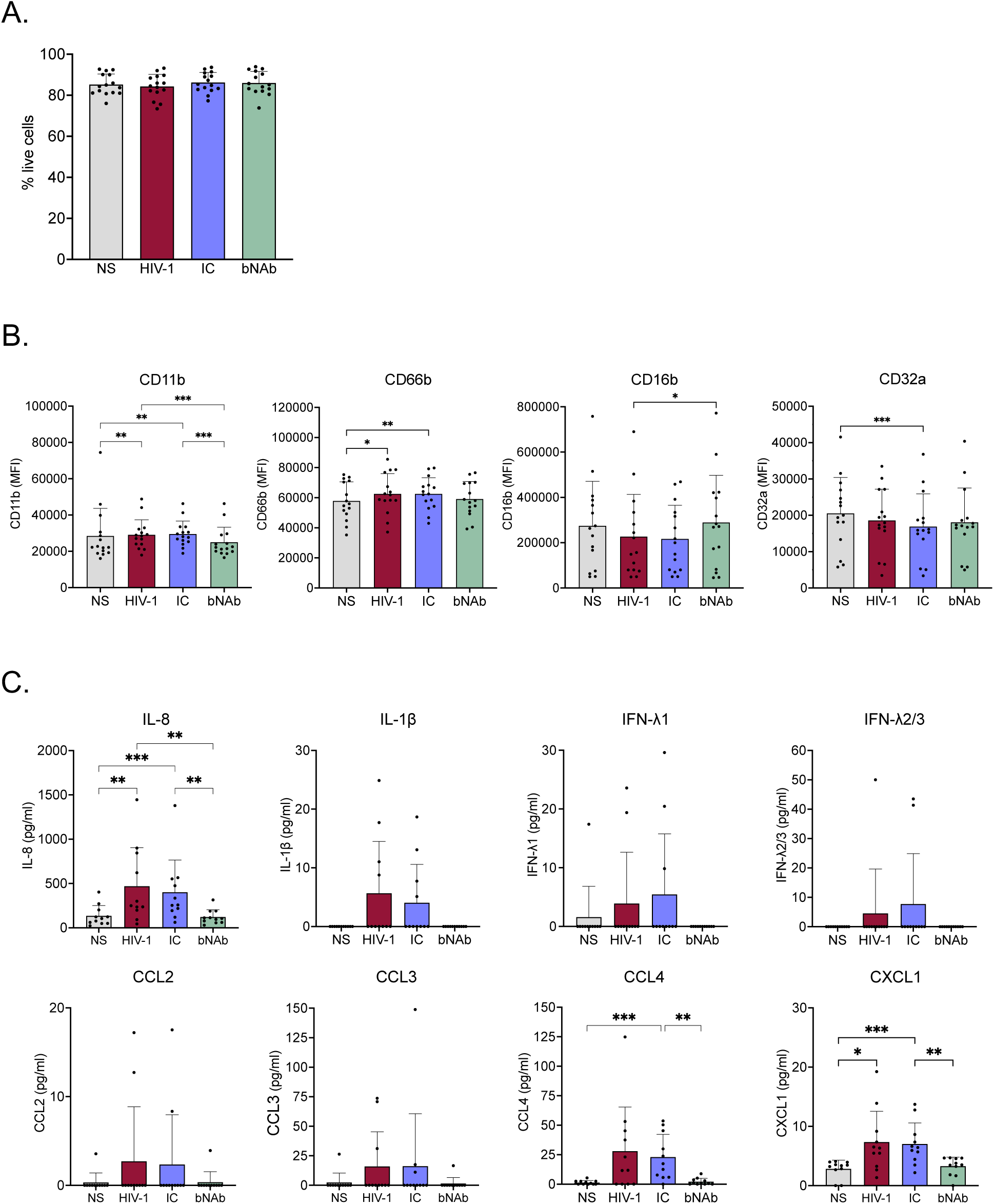
Activation of human neutrophils stimulated *in vitro* by HIV-1 virions, ICs or free bNAb. Neutrophils from HD were activated overnight by HIV-1 virions, ICs or free bNAb. Activated neutrophils and corresponding cell culture supernatants were collected to evaluate neutrophil viability and phenotypic activation via flow cytometry, as well as to analyze their secretome using bead-based immunoassays. (A-B) Viability (A) and phenotypic activation (B) assessed by monitoring cell surface markers. (C) Cytokines and chemokines secretion profile was performed as in Figure 1 and is represented for each cytokine/chemokine detected. Values are mean +/− SD of at least 15 (A-B) and 11 (C) independent measurements. Several cytokines/chemokines were detected only in a fraction of neutrophil samples assayed. Significance was assigned as follows: ∗p < 0.05, ∗∗p < 0.01 and ∗∗∗p < 0.001, NS: non-stimulated.

### HIV-1 activation of neutrophils induces IL-8, CXCL1, and CCL4 secretion

In parallel, we assessed the functional activation of neutrophils isolated from HD by HIV-1 virions, free or in the form of ICs. Free 10-1074 bNAb was used as control and showed no effect on either the expression of cell surface markers or the secretome of neutrophils. Upon exposure to HIV-1 and ICs, neutrophils displayed a measurable activation profile, characterized by the secretion of specific inflammatory mediators, although to a lesser extent as compared to TLR mimicking viral stimuli (TLR7/8). Notably, IL-8, CXCL1, and CCL4 were significantly upregulated compared to unstimulated cells (**Figure 3C**), indicating a preserved capacity to initiate chemotactic and antiviral responses. Additional cytokines and chemokines, including IL-1β, IFN-λ1, IFN-λ2/3, CCL2, and CCL3, were also detectable in a subset of donors, reflecting a variable but inducible immune responsiveness. Importantly, the overall secretome profiles elicited by HIV-1 and ICs were comparable, supporting the notion that neutrophils retain functional responsiveness to both types of stimuli. Collectively, these results highlight a selective and functionally preserved proinflammatory profile of neutrophils upon HIV-1 encounter, characterized by a balanced and targeted chemokine release without excessive activation, including in the context of antibody therapy.

### Neutrophils from PLWH exhibit increased release of specific cytokines and chemokines upon stimulation with danger signals associated with HIV-1 infection

Neutrophil effector functions are known to be impaired during HIV-1 infection (5,6). Notably, the restoration of these functions with ART has also been evaluated (31). However, the impact of HIV-1 infection on neutrophil cytokine and chemokine secretion profiles remains underexplored, with most studies focusing on individuals at the AIDS stage (32) rather than ART-treated PLWH. Thus, it is important to investigate how HIV-1 infection and ART treatment affect their immunomodulatory potential. To this end, we evaluated the activation of neutrophils isolated from ART-treated PLWH (**Supplementary Figure 3A–D**) in response to TLR agonists, pro-inflammatory cytokines, HIV-1, or ICs, and compared their responses to neutrophils isolated from HD, including newly collected samples. Importantly, no significant differences were observed in neutrophil frequency in the blood of PLWH compared to HD (**Supplementary Figure 3E**) regardless of the time since diagnosis (**Supplementary Figure 3F**). Our findings reveal that neutrophils from ART-treated PLWH not only retain their capacity to secrete cytokines and chemokines in response to various danger signals —including bacterial- and virus-like stimuli and pro-inflammatory cytokines— but also exhibit an enhanced release of specific mediators compared to HD (**Figure 4**). Notably, upon TLR4 stimulation, neutrophils from PLWH showed increased secretion of CXCL1 and CCL3, while TLR7/8 activation led to elevated levels of CCL3 and IFN-γ (**Figure 4A, 4B**). Interestingly, this new set of samples confirmed that TLR4-activated neutrophils can secrete IFN-λ2/3, as observed in the majority of samples from both HD and PLWH (**Figure 4B**). Pro-inflammatory cytokines differentially modulated the secretion profiles of neutrophils from PLWH compared to those from HD (**Figure 4C, 4D**). Specifically, TNF-α stimulation enhanced CCL2 secretion but reduced IFN-λ1 and IFN-λ2/3production by PLWH neutrophils. Furthermore, IFN-γ stimulation induced the secretion of CCL3 and CCL4 exclusively in neutrophils from PLWH (**Figure 4B**, 4D). This heightened and selective responsiveness to TLR agonists and pro-inflammatory cytokines suggests a functionally primed state, likely resulting from chronic immune activation or low-grade inflammation associated with HIV-1 infection. In contrast, neutrophils from PLWH had similar secretion profiles than those from HD upon stimulation with free HIV-1 virions or in the form of ICs, both qualitatively (type of cytokine/chemokine) and quantitatively (amount secreted), with the exception of a trend toward reduced CCL4 production in response to HIV-1 and IC stimulation (**Supplementary Figure 4)**. These results highlight a distinct neutrophil secretory profile in ART-treated PLWH, particularly under certain stimulatory conditions and/or in specific donor samples. This secretory profile is characterized by both preserved and selectively (stimulus-dependent) heightened immunomodulatory responses. This nuanced cytokine and chemokine landscape suggests a complex yet functionally relevant adaptation of neutrophils during HIV-1 infection, even under effective ART, with potential implications for host defense and inflammation regulation.

**Figure 4.**
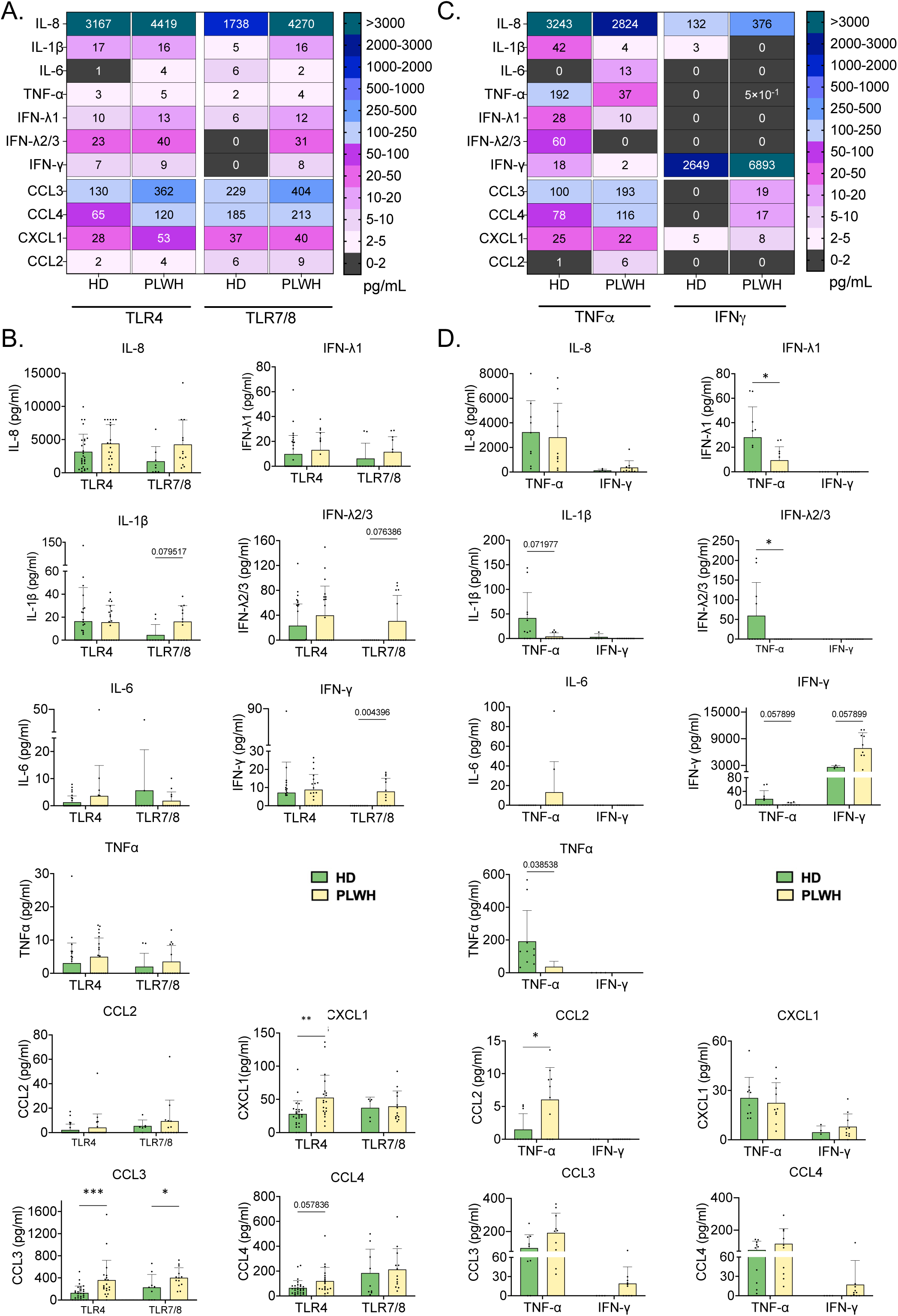
Activation of neutrophils isolated from HD or PLWH stimulated *in vitro* by TLR agonists LPS (TLR4) and R848 (TLR7/8) or inflammatory cytokines (TNF-α and IFN-γ). Neutrophils from HD were activated overnight by TLR agonists or pro-inflammatory cytokines. Cell culture supernatants of activated neutrophils were collected to analyze their secretome using bead-based immunoassays, as for Figure 1. (A-B) Cytokines and chemokines secretion profile upon activation by TLR agonists, represented as a heatmap (A) and histograms (B) for each cytokine/chemokine assessed. (C-D) Cytokines and chemokines secretion profile upon activation by inflammatory cytokines, represented as a heatmap (C) and histograms (D) for each cytokine/chemokine assessed. Values are mean +/− SD of at least 9 independent measurements. Several cytokines and chemokines were only detected in a fraction of samples assayed.

### Altered FcγR expression dynamics in neutrophils of PLWH

We analyzed FcγR expression on neutrophils from PLWH in response to various danger signals associated with HIV-1 infection, which is particularly relevant in the context of bNAb-based therapies. At baseline, neutrophils from PLWH exhibited significantly higher surface expression of CD16b compared to those from HD (**Supplementary Figure 5A**), while CD32 levels showed no significant difference between the two groups (**Supplementary Figure 5B**). Following stimulation with TLR agonists, pro-inflammatory cytokines, HIV-1 virions, or ICs, overall surface expression levels of CD16b and CD32a were comparable between the two groups (**Supplementary Figure 5A-B**). However, when assessing the modulation of FcγR expression relative to baseline levels, distinct differences emerged. Upon TLR4 activation, neutrophils from PLWH exhibited a significantly greater reduction in surface CD16b and CD32a expression compared to HD (**Figure 5A**). In contrast, TLR7/8 stimulation did not induce significant differences in receptor downmodulation between the groups (**Figure 5A**). Similarly to TLR4, TNF-α stimulation caused a more pronounced decrease in CD16b and CD32a expression on neutrophils from PLWH, while IFN-γ selectively reduced CD16b expression (**Figure 5B**). Stimulation with free HIV-1 virions or ICs did not result in any notable differences in FcγR modulation between PLWH and HD (**Figure 5C**). In summary, neutrophils from PLWH exhibit elevated baseline levels of CD16b and show enhanced downmodulation of CD16b and CD32a in response to TLR4 and inflammatory cytokine stimulation. However, this heightened FcγR downregulation was not observed following exposure to viral stimuli such as TLR7/8 agonists or HIV-1 virions. These findings reveal distinct FcγR regulatory dynamics in neutrophils from PLWH, shaped by both the inflammatory environment and the nature of the activating stimulus.

**Figure 5.**
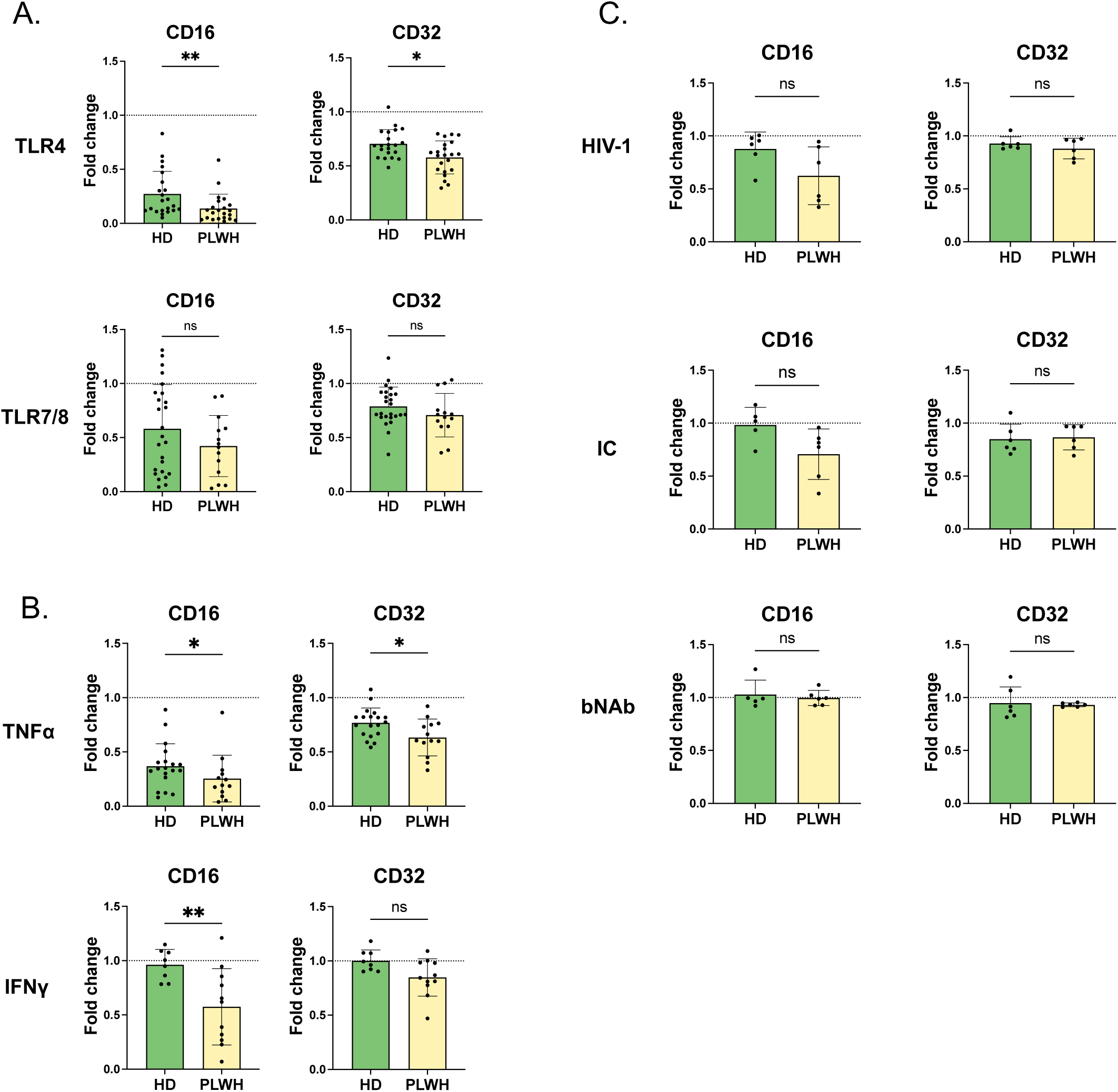
Modulation of the expression of FcγR on neutrophils isolated from HD or PLWH stimulated *in vitro* by TLR agonists, inflammatory cytokines, HIV-1 virions, ICs or free bNAb. (A-C) Modulation of the cell surface levels of CD16 and CD32 upon activation by TLR agonists (A), cytokines (B), and HIV-1 virions, ICs or free bNAb (C). Fold change is measured versus unstimulated cells (dashed lines). It is represented as the ratio mean fluorescence intensity (MFI) stimulated cells/MFI unstimulated cells. Values are mean +/− SD of at least 14 (A), 8 (B) and 5 (C) independent measurements. Significance was assigned as follows: ∗p < 0.05 and ∗∗p < 0.01

### Neutrophils from PLWH display enhanced expression of activating FcγRs, CD63, CXCR4 and PD-L1 as well as reduced expression of CD11b and CD49d

To further evaluate how HIV-1 infection impacts neutrophil properties, we conducted a phenotypic analysis of neutrophils isolated from PLWH and HD. We measured the expression of 18 cell surface markers under basal conditions, focusing on markers relevant to HIV-1 infection (CXCR4), neutrophil activation (CD11b, CD66b, CD63, CD62L), maturity (CD10), immunosuppressive properties (PD-L1), and Fc receptors (CD16, CD32, CD64, CD89), among others. Notably, we confirmed the elevated expression of the activating FcγR CD16b and observed a significantly higher expression of the inducible activating FcγR CD64 in neutrophils from PLWH. Additionally, we identified increased expression of the degranulation marker CD63, as well as increased levels of CXCR4 and PD-L1. Conversely, decreased expression of CD49d was detected in PLWH neutrophils (**Figure 6**). These findings demonstrate distinct phenotypic differences in neutrophils from PLWH compared to HD, which may underlay altered functional properties, consistent with the heightened responsiveness to inflammatory stimuli observed in this study.

**Figure 6.**
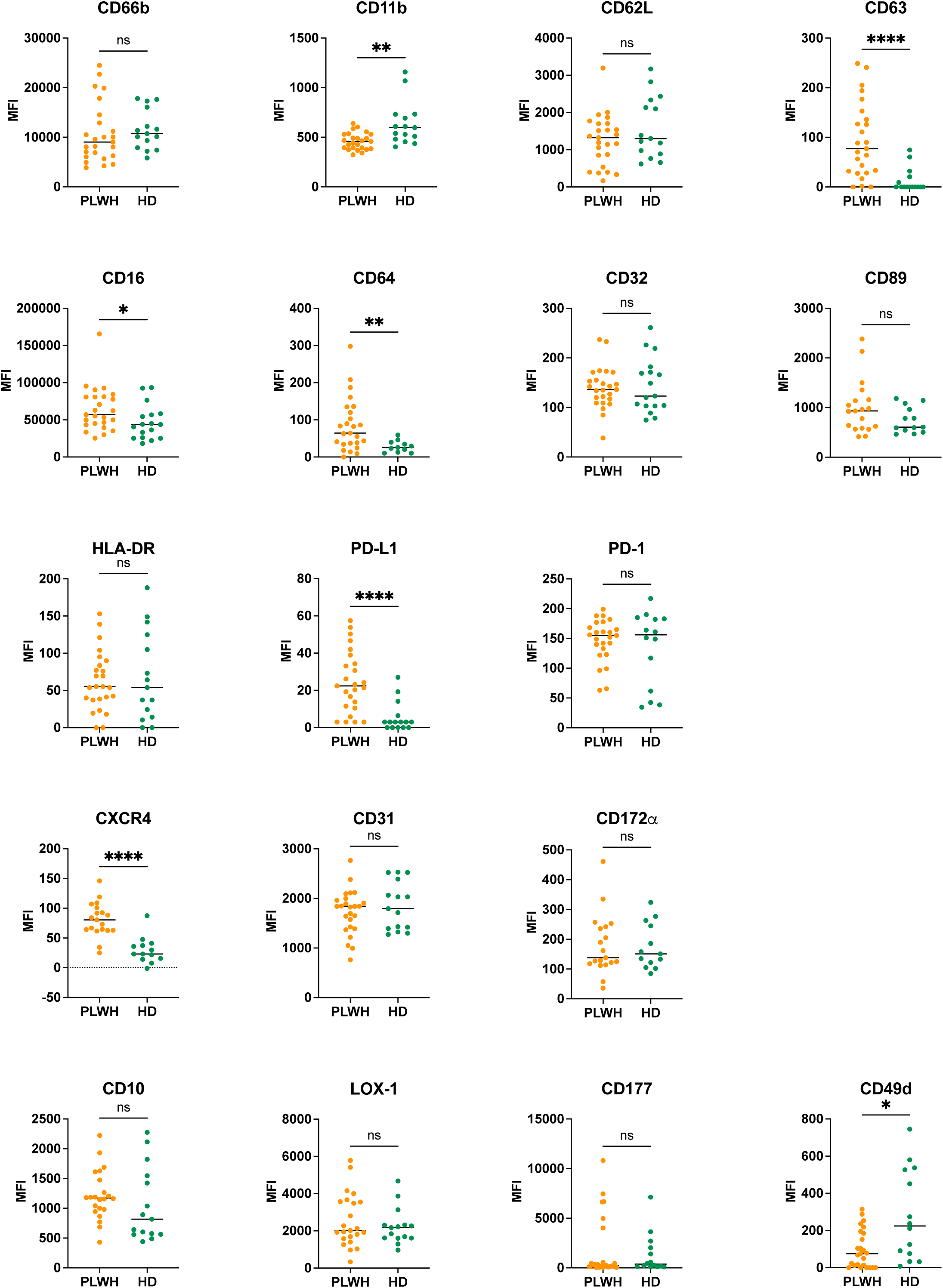
Phenotypic characterization of neutrophils isolated from PLWH and HD blood samples. Neutrophils purified from whole blood of HD and PLWH were immunophenotyped by assessing the expression of 18 cell surface markers using multiparametric flow cytomety. Data include at least 19 PLWH blood samples and 12 HD samples. Statistical analyses were done using unpaired Mann-Whitney test. Significance was assessed as follows: ∗p < 0.05, ∗∗p < 0.01, and ∗∗∗∗p < 0.0001.

## Discussion

We previously demonstrated in a mouse model of retroviral infection the crucial immunomodulatory role of neutrophils in inducing protective (21,22). Here, we provide new insights into the immunomodulatory properties of neutrophils in the context of HIV-1 infection and antiviral antibody therapy, providing a detailed analysis of their secretome and immune receptors expression in a stimulus-specific manner, comparing HD and PLWH. Our work shows that neutrophil from ART-treated PLWH retain their capacity to secrete cytokines and chemokines in response to danger signals and, upon certain stimuli, exhibit enhanced secretion of specific soluble mediators compared to neutrophils from HD, suggesting the effect of HIV-1 infection in neutrophil properties. In addition, notable differences in phenotypic features distinguish them from neutrophils of HD origin. Noteworthy, most studies on immunomodulatory functions of neutrophils have mostly focused on bacterial activation (i.e. TLR4 activation), while the effect of virus-dependent activation is understudied, often relying on TLR-based surrogate models. Our findings fill this knowledge gap by revealing how HIV-1 and/or HIV-ICs influence the immunomodulatory functions of neutrophils, and suggest that neutrophils in PLWH maintain functional plasticity and may play a role in shaping immune responses.

Our data show that both TLR4 and TLR7/8 agonists modulate FcγR expression on neutrophils. Specifically, the TLR4 agonist significantly downregulates CD16b and CD32a, while the TLR7/8 agonist selectively reduces CD32a expression, as previously reported (33). These findings suggest that bacterial signals exert a more pronounced regulatory effect on FcγR expression—particularly on CD16b—compared to viral stimuli. Consistently, exposure to free HIV-1 or ICs has minimal impact on CD16b levels. Additionally, TNF-α, but not IFN-γ, markedly decreases CD16b expression and modestly reduces CD32a. The reduction in FcγR cell surface levels can occur through various mechanisms, including receptor internalization and shedding of the extracellular portion of the FcγR (33,34). Worthy of note, the decrease in FcγR expression following cell activation may serve as a regulatory mechanism, modulating receptor availability on the cell surface and influencing FcγR-mediated cell activation. This stimulus-dependent and pathology-dependent modulation of FcγRs (i.e. increased CD16b and CD64 basal level observed in PLWH neutrophils) might have biological consequences and should be considered in the context of HIV-1 infection and bNAb-based immunotherapies, whether alone or in combinatorial therapeutic approaches involving TLR agonists (14,35–39) as it could influence their therapeutic efficacy. This might also be relevant to consider it in the context of bacterial co-infections in PLWH (40).

Our work also allowed to dissect at the protein level the chemokine/cytokine secretion profile of neutrophils in a stimulus-dependent manner under both physiological (HD) and pathological (PLWH) conditions. This is important to point out as multiple studies assessing the secretion profile of neutrophils upon different stimuli were based on transcriptomic approaches instead of *bona fide* cytokine/chemokine secretion. Consistent with prior findings, we have shown that neutrophils from HD secrete IL-8, CXCL1, CCL3, and CCL4 upon LPS, R848, or TNF-α activation (25–27,29), while IFN-γ only induced a low secretion of IL-8. Additionally, our work finely describes the secretome of HIV-1-and IC-activated neutrophils, thus far unreported. The secreted cytokines and chemokines, IL-8, CXCL1 and CCL4, play roles in neutrophil activation and recruitment, as well as in recruiting NK cells, monocytes, and other immune cells to sites of inflammation or tissue damage (41). These effects could either amplify inflammation through the recruitment of myeloid cells or enhance viral control through the activation of diverse effector cells. Interestingly, the secretion of different types IFN-λ by neutrophils upon LPS, TNF-α and HIV-1 stimulation, though observed in only a subset of samples, is a novel finding. Worthy of note, IFN-λ1 and IFN-λ2/3 secretion was significatively reduced in TNF-α-activated neutrophils from PLWH, suggesting that the inflammatory environment associated with HIV-1 infection might downregulate IFN-λ secretion by neutrophils. This family of cytokines have been shown to modulate antiviral immune response (42), but its role in HIV-1 infection is still unclear. Further research is needed to confirm IFN-λ secretion by neutrophils under different stimuli and to understand its biological implications in neutrophil function and HIV-1 infection. In this regard, it has been shown that IFN-λ (i) diminishes the production of ROS and degranulation in neutrophils (43) and (ii) IFN-λ3 inhibits HIV-1 infection of macrophages (44).

Globally, our findings underline that neutrophils isolated from ART-treated PLWH exhibit similar responses to HIV-1 and ICs as neutrophils from HD but exhibit heightened reactivity to TLR agonists and pro-inflammatory cytokines. Notably, the neutrophil secretome in the context of long-term ART has remained largely uncharacterized until now. Our data contrast with earlier reports describing impaired cytokine secretion by LPS-stimulated neutrophils from PLWH who had progressed to AIDS and had been on ART for less than two years (32). These discrepancies suggest that in advanced disease stages, neutrophil responsiveness to pathogen-associated signals may be compromised, potentially contributing to disease progression. In contrast, our results highlight that ART not only preserves neutrophil effector functions (6,45), but also their cytokine/chemokine secretion capacity. Additionally, our work sheds new light on the phenotypic characteristics of neutrophils in PLWH. First, we observed an increased basal expression of two activating FcγRs, CD16b and CD64, while CD32a levels remained unchanged. Notably, CD64 expression on neutrophils is inducible by IFN-γ (46,47) (**Supplementary Figure 1**), supporting a model in which IFN-γ–mediated conditioning alters neutrophil properties in PLWH. This IFN-γ–mediated conditioning is further consistent with the elevated expression of PD-L1 observed on neutrophils from PLWH. The upregulation of CD16b and CD64 could potentially enhance neutrophil binding to viral ICs. However, this increase did not correlate with enhanced cytokine or chemokine secretion upon IC stimulation, suggesting that these two FcγR are not the primary drivers of cytokine/chemokine induction in response to HIV-ICs. Instead, this points toward a potential role for CD32a—which remained unchanged—in mediating these Fc-dependent responses. This interpretation aligns with the known functional specialization of FcγRs: CD32a is a potent activator of phagocytosis and inflammatory signaling, while CD16b mainly promotes β1 integrin activation, contributing to a pro-adhesive phenotype and the formation of NETs (15). Further investigations are required to determine the potential FcγR-dependent enhancement of cytokines/chemokine by HIV-ICs as well as the FcγRs involved. Additionally, the upregulation of FcγRs could play a role in clearing infected cells through ADCC (48,49) but also in promoting inflammatory immune responses such as reactive oxygen species (ROS) production (50) and NET formation (51). These phenomena can be key in antiviral immune defense but, if exacerbated, they can induce pathological inflammation. In this regard, upregulation of these FcγRs could point to a role for neutrophils in promoting/maintaining chronic inflammation in PLWH. Second, we observed increased expression of CD63, a marker of degranulation that is associated with the release of granules containing enzymes, antimicrobial peptides, and other effector molecules (52,53) suggesting that neutrophils from PLWH might be actively degranulating, possibly in response to infection or inflammation. Third, our results also show an enhanced expression of CXCR4 and PD-L1. Interestingly, the upregulation of these two cell surface molecules has recently been associated with neutrophil aging in PLWH (54). However, in contrast to our study, the assessment of CXCR4 and PD-L1 expression was made using samples from ART-naive PLWH. Elevated CXCR4 expression may enhance chemotaxis towards inflammatory sites, potentially aiding immune surveillance but also raising concerns about facilitating HIV-1 dissemination (55). Worthy of note, enhanced expression of CXCR4 was observed at any clinical stage of PLWH (**Supplementary Figure 6**). The concurrent decrease in CD49d expression, an integrin subunit essential for adhesion and migration, could indicate altered migratory behaviors, favoring CXCR4-mediated pathways. Globally, the enhanced expression of FcγRs, CD63, CXCR4 as well as the decreased expression of CD49d suggests that neutrophils from PLWH show a state of heightened activation, readiness, and targeted migration of neutrophils. It may suggest an adaptive response for rapid pathogen clearance and active participation in acute inflammatory responses. However, it could also imply a risk of pathological inflammation or tissue damage if not properly controlled. Finally, we confirmed the upregulation of PD-L1 in neutrophils from PLWH, consistent with previous studies describing its expression across various neutrophil subsets (54,56,57) and irrespective of ART (54,56) in PLWH. PD-L1 expression has been associated with T-cell immunosuppressive effects that can be counteracted by blocking the PD-1/PD-L1 axis (54,56), suggesting a potential immunosuppressive role for neutrophils in PLWH.

In conclusion, our work dissects the phenotypic and functional properties of PLWH neutrophils. While these cells retain their capacity to secrete cytokines and chemokines, they also exhibit immunosuppressive features, such as PD-L1 expression. Our findings also provide new insights on the modulation of FcγR expression by pro-inflammatory cytokines and pathogen-associated danger signals relevant to HIV-1 infection—factors that should be considered in the context of antibody-based therapies. In addition to providing a comprehensive analysis of neutrophil immunomodulatory properties and FcγR dynamics in both health and HIV-1 infection, these findings have important implications for the development of tailored antiviral therapies for PLWH, particularly those involving bNAbs, either alone or in combination with TLR agonists. Moreover, these insights could inform future combinatorial approaches designed to enhance immune responses by targeting neutrophil-mediated regulatory mechanisms—extending beyond bNAb-based interventions— to include strategies such as PD-1/PD-L1 blockade to counteract immunosuppression (54,56) or advanced therapies aimed at limiting potential neutrophil-driven pathological inflammation.

## Material & Methods

### Human blood samples

Blood samples from anonymized HD were provided by the French establishment for blood donations (Etablissement Français du Sang, EFS, Montpellier, France) according to the agreement between this establishment and INSERM (21PLER2018-0069). EFS provided 5mL of fresh whole blood in EDTA for each healthy donor. The use of blood samples from anonymized PLWH under ART was approved by the Centre for Biological Resources (DRI#305-2019-NC and amendment DRI_2022-69-SR). The samples were collected and provided by the Virology laboratory of the Centre Hospitalier Universitaire (CHU) Lapeyronie of Montpellier (France). Median age was 57 years old ([22;76]) for 31 PLWH donors (19M; 13F). All donors were under successful ART with no detectable viral load. 2 to 3mL of whole blood in EDTA were provided for each PLWH donors.

### Human neutrophil purification

Whole blood sample was diluted to three times its volume with a solution of PBS 1X (Gibco, cat #10010-015) supplemented with EDTA at 0,2 mM (Invitrogen, cat #15575-038) and loaded on Histopaque-1119 (Sigma-Aldrich, cat #11191). Density gradient was performed by 800g centrifugation with slow acceleration and no brake (acceleration 4, brake 0) for 20 minutes. Granulocytes phase was collected, washed with PBS containing EDTA (0,2 mM) and 2% FBS (FBS, Eurobio, REF#CVFSVF00-01) and neutrophils were purified with CD15-positive selection kit (Stemcell technologies, CD15+ positive selection REF#18651) as previously described in (57). Purity assessed by flow cytometry was > 98%.

### Cell culture

Purified neutrophils were plated at a concentration of 2 million cells/mL in 96 U-bottom wells plate (3 x 10^5^ /well) and cultured in RPMI 1640 supplemented with GlutaMAX^TM^ medium (Gibco, cat #61870-010), 10% fetal bovine serum (FBS, Eurobio, REF#CVFSVF00-01), 1% penicillin/streptomycin (Gibco, cat #15140-122) and 10 ng/mL of G-CSF (R&D, cat #214-CS-025).

### In vitro cell stimulation

Cells were stimulated overnight using TLR agonist, inflammatory cytokines or viral determinants under the following conditions: (i) TLR4 agonist (LPS; Sigma-Aldrich, cat #L4524) or TLR7/8 agonist (R848; Invivogen, cat #tlrl-r848) at 0,5 μg/mL each; (ii) IFN-γ (R&D, cat #285-IF-100) at 50ng/mL or TNF-α (Peprotech, cat #AF-300-01A) at 1ng/mL; (iii) free HIV-1 virions (Ad8 strain, 3.10^4^ viral particles) or ICs made with HIV-1 Ad8 and the 10-1074 bNAb (as described below) or the bNAb alone (at 1μg/mL). Dose-response experiments were previously conducted to determine the optimal stimulation conditions.

### Virus production, purification and titration

2.5 million human embryonic kidney cells (293THEK cell) were seeded in 10 ml of growth media 1 day before transfection. At 50-70% confluence, cells were transfected with calcium phosphate precipitate method, with 10 µg total quantity of HIV-1 molecular clone expressing plasmids, as in (58). At 12 hours post transfection, cells were washed and fresh media supplemented with 20mM Hepes was added on the cells. Cell culture supernatant containing viral particles was collected 36 hours post-transfection, pooled and filtrated through 0.45 µm and then purified by ultracentrifugation on a cushion of 25% sucrose in TNE buffer (10 mM Tris-HCl [pH 7.4], 100 mM NaCl, 1 mM EDTA) at 100000xg, for 1 hour 30 minutes at 4°C, in a SW32Ti Beckman Coulter rotor. Dry pellet was resuspended with RPMI without serum, at 4°C overnight. The viral particles were quantified using Videodrop system (Myriade, France).

### Recombinant HIV-1 bNAb production

Recombinant human HIV-1 IgG1 antibodies, anti-CD4bs 3BNC117 (59) and anti-V3-glycan 10-1074 (60), were produced by co-transfection of Freestyle 293-F cells (Thermo Fisher Scientific) using the polyethyleneimine (PEI)-based precipitation method as previously described (61), and purified by affinity chromatography using protein G Sepharose 4 fast flow beads according to the manufacturer’s instructions (Cytiva). Purified IgG antibodies were dialyzed against PBS using Slide-A-Lyzer Cassettes (10K MWCO; Thermo Fisher Scientific), and quantified using NanoDrop™ ONE instrument (Thermo Fisher Scientific). Both bNAbs are of IgG1 isotype and were used at a concentration of 1μg/mL to form ICs.

### Immune complexe formation

Viruses (3.10^4^ viral particles) and bNAb (1μg/mL) were diluted in culture medium and mixed together on a 1,5mL Eppendorf, then mixed on a rotative agitator for 20 minutes. ICs were gently plated with cells in 96 U-bottom wells plate.

### Flow cytometry immunophenotyping of neutrophils

For monitoring phenotypic activation, cells were stained at room temperature with a viability dye (ZombieRed^TM^, Biolegend #423109) for 15 minutes then washed and stained for 20 minutes at room temperature with fluorochrome-conjugated antibodies. Antibodies for flow cytometry were PerCPCy5-5-CD15 (HI98, #301922), AF488-CD11b (ICRF44, #301317), AF647-CD66b (G10F5, #305110), PC7-CD16a/b (3G8, #302016), PE-CD32a/b (FUN-2, #303206) and PE-CD64 (10.1, #305008) from BioLegend. Finally, cells were washed and resuspended with PBS1X. Data acquisition was performed on a Novocyte (Agilent Technologies) flow cytometer.

For immunophenotyping of HD’s and PLWH’s neutrophils the following antibodies were used: PerCPCy5-5-CD15 (HI98, #301922), PE-LOX-1 (15C4, #358604), BV510-CD11b (ICRF44, #301334), AF647-CD66b (G10F5, #305110), BV605-CD62L (DREG-52, #304834), BV650-HLA-DR (L243, #307650), BV711-CD63 (H5C6, #353042), PC7-CD16a/b (3G8, #302016), PE-CF594-CD64 (10.1, #565389), APC-Fire750-CD32a/b (FUN-2, #303220), APC-CD177 (MEM-166, #315808), BV605-CD49d (9F10, #304324), BV711-CD31 (WM59, #303136), PE-CF594-CD10 (HI10a, #312227), PE-CD89 (A59, #555686), PC7-CXCR4 (12G5, #306514) from BioLegend. Additionally, PE-PD-1 (3.13, Miltenyi #130-177-384), PC7-PD-L1 (MIH1, BD #558017) and APC-CD172*α* (15-414, eBiosciences #17-1729-42). The viability staining was made by resuspending the cells with 1mL DAPI (Biolegend #422801) diluted at 3*µ*M with an incubation of 15 minutes at room temperature. After the washing of fluorescent antibodies, cells were fixed with 100 μL PFA4% for 10 minutes at room temperature. Finally, cells were washed and resuspended with PBS1X. Data collection was carried out using a BD LSRFortessa flow cytometer from BD Bioscience.

Forward scatter area and forward scatter height as well as side scatter area and height were used to remove doublets from flow cytometry analyses. Gating is done on live cells as assessed by viability dye and on CD15-positive cells. Data were analyzed using FlowJo software version 10.5.3 (TreeStar).

### Cytokines and chemokines secretion quantification

Secretion of soluble cytokines and chemokines were assayed from cell-free culture supernatants of *in vitro* stimulated neutrophils using bead-based immunoassays (LEGENDplex^TM^, Biolegend). Two assays were used allowing the quantification of IL-1β, Il-6, TNF-α, IP-10 (CXCL10), IFN-λ1, IL-8 (CXCL8), IL-12p70, IFN-α2, IFN-λ2/3, GM-CSF, IFN-β, IL-10, IFN-γ (LegendPlex Human anti-virus response panel, Biolegend #740390) and CCL11, CCL17, CCL2, CCL5, CCL3, CXCL9, CXCL5, CCL20, CXCL1, CXCL11, CCL4 (LegendPlex Human proinflammatory chemokines panel, Biolegend #740985). Immunoassays plates were read on a Novocyte (Agilent Technologies) flow cytometer and data were analyzed with the appropriate Biolegend LEGENDplex^TM^ software (https://legendplex.qognit.com).

### Surface Plasmon resonance (SPR)

Affinity of bNAbs for FcγRs was assessed by SPR experiments performed on a Biacore T200 (GE Healthcare). SPR experiments were performed on a T200 apparatus at 25°C in PBS containing 0,05 % P20 surfactant (Cytiva). Anti-histidine antibody (R&D Systems) was covalently immobilized on a CM5-S sensor chip flowcell (Fc2) by amine coupling according to the manufacturer’s instructions (Cytiva). A control reference surface (flowcell Fc1) was prepared using the same chemical treatment but without anti-His antibody. All kinetic measurements in Fc1 and Fc2 were performed by single-cycle titration at 100μl/min. Each human FcγR (R&D Systems) was captured on immobilized anti-His antibodies at 100-200 RU level. Five increasing concentrations (3,6, 11, 33, 100, 300 nM) of antibody were injected (injection time = 60s) at 100µl/min on captured receptors. After a dissociation step of 600s in running buffer, sensor surfaces were regenerated using 10μl of glycine-HCl pH1.5. All the sensorgrams were corrected by subtracting the low signal from the control reference surface and buffer blank injections. Kinetic parameters were evaluated from the sensorgrams using a two-states or a steady-state models from the T200 evaluation software.

### Statistics

Statistical analyses were performed using Prism software version 9.5.1 (GraphPad). Simple group comparisons were performed using the Wilcoxon signed rank test for paired data or the Mann–Whitney test for unpaired data. Multiple group comparisons were performed using Friedmann test for paired data or using Kruskal-Wallis test for unpaired data with additional multiple comparisons Dunn test. Significance was assigned as follows: ∗p < 0.05, ∗∗p < 0.01, ∗∗∗p < 0.001, and ∗∗∗∗p < 0.0001.

## Acknowledgements

This work was supported by grants from Sidaction (A014-2-AEQ-08-01; BI25-1-02278, 20-1-AEQ-12652), ANRS (France Recherche Nord&sud Sida-hiv Hépatites; ECTZ46143; ECTZ47079, ECTZ204786), and INSERM state funding granted to M. Naranjo-Gomez (U1183NAR). This work was also supported by the consortium ACT4COVIDCellnex (220110FF, RAK200019FFA). R. Dibsy was a recipient of a Sidaction PhD fellowship. S. Marsile-Medun, M. Naranjo-Gomez, Manon Souchard, Daouda Abba Moussa, Martine Pugnière and M. Pelegrin are members of the “MabImprove Labex”, a public grant overseen by the French National Research Agency (ANR) as part of the “Investments for the future” program (reference: ANR-10-LABX-53-01) that also supported this work. We thank the imaging facility MRI, which is part of the UAR BioCampus Montpellier and a member of the national infrastructure France-BioImaging, supported by the French National Research Agency (ANR-10-INBS-04, “Investments for the future”). We thank the BSL3 research laboratory CEMIPAI (CNRS UAR3725) for providing help and advice, and the Centre de Ressources Biologiques (CRB) for providing blood samples from PLWH. We thank the Montpellier Proteomics Platform (PPM-PP2I, BioCampus Montpellier) facilities where the SPR experiments were carried out. We are grateful to Dr. Paris for critical reading of the manuscript.

## Authors contribution

Soledad Marsile-Medun (SM-M), Mar Naranjo-Gomez (MN-G) and Mireia Pelegrin (MP) defined the research program; SM-M, Manon Souchard (MS), Daouda Abba Mousssa (DAM), Giang Ngo (GN), Martine Pugnière (MPu,) and MN-G, performed the experiments; Valerie Lorin (VL) and Hugo Mouquet (HM) provided advice on and generated bNAbs; Elisa Reynaud (ER) and Edouard Tuaillon (ET) managed and provided PLWH blood samples. Rayane Dibsy (RD) and Delphine Muriaux (DM) provided advice on and performed HIV-1 production, purification and quantification. SM-M, MS, MN-G and MP carried out the data analyses. SM-M, MN-G and MP wrote the manuscript. Grants to SM-M, MN-G and MP funded the study. All authors discussed, commented on and approved the manuscript in its final form.

