## Supplemental Figures for "Comprehensive Analysis of Neutrophil Immunomodulatory Properties and FcR Dynamics in Health and HIV-1 Infection and Therapy"

### Supplementary Figures

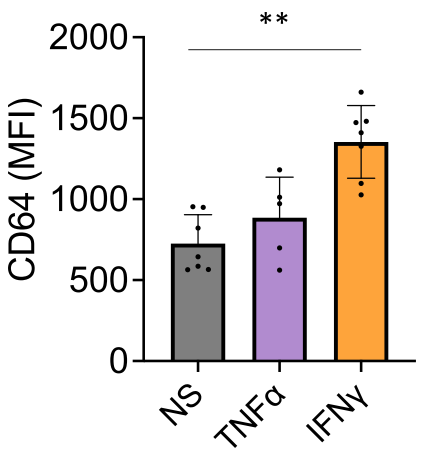


**Supplementary Figure 1. Assessment of cell surface expression of FcγRI (CD64) on neutrophils activated with TNFα at 1 ng/ml and IFN-γ at 50 ng/ml.** Values are mean +/- SD of 5 (TNFα) and 7 (IFNγ) independent measurements. Significance was assigned as follows: **p < 0.01.

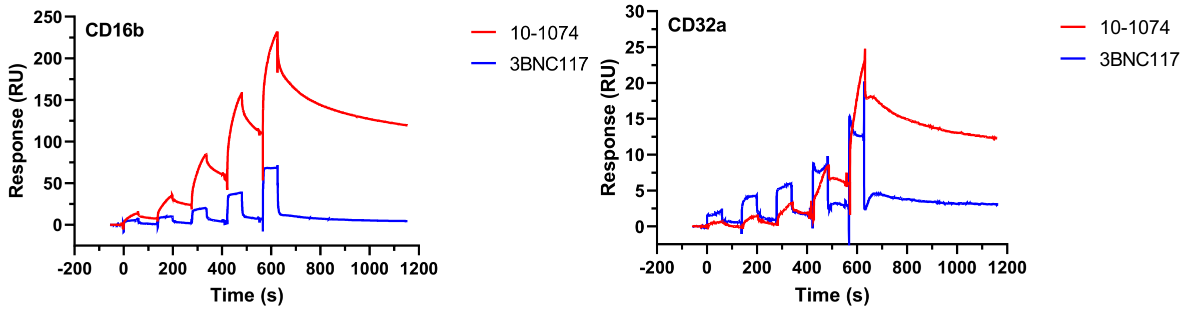


| Sample | Rmax (RU) | Chi² (RU²) | Ligand | Ligand Level (RU) | KD | Fit |
|---|---|---|---|---|---|---|
| **3BNC117** | 109 | 1,14 | CD16b | 200 | 2,1 µM | Steady State |
| **10-1074** | 208 | 8,8 | CD16b | 200 | 44 nM | Two State Reaction |
| **3BNC117** | 13,7 | 0,6 | CD32a | 140 | 1,1 µM | Steady State |
| **10-1074** | 77,9 | 0,08 | CD32a | 140 | 0,36 µM | Two State Reaction |

**Supplementary Figure 2. Surface plasmon resonance (SPR) analysis of FcγRIIa (CD32a) and FcγRIIIb (CD16b) binding to 3BNC117 and 10-1074 bNAbs.**
Kinetic measurements were performed by single-cycle titration on a T200 apparatus.

**A**

| Sample ID | Age | Gender | Years since: Diagnostic | Years since: Indetectable viral load | ART regimen | CD4 count (mm³) | CD4/CD8 ratio | Viral load (cp/mL) |
|---|---|---|---|---|---|---|---|---|
| ***21-271-2023*** | 50 | M | 21 | >4 | DOL+ABC+3TC | 446 | 0,9 | 0 |
| ***21-271-1719*** | 52 | M | 2 | 2 | TAF+FTC+BIC | 719 | 1 | 0 |
| ***21-280-2233*** | 54 | F | 34 | >4 | RAL+DRV+RTV | 1112 | 1 | 0 |
| ***21-280-2392*** | 52 | M | 32 | >4 | DOL+ABC+3TC | 1175 | 1,2 | <20 |
| ***21-294-1519*** | 59 | M | 7 | >4 | TAF+FTC+EVG+cobi | 567 | 1 | 0 |
| ***21-294-2073*** | 57 | M | 19 | >4 | LPV/r | 734 | 2,3 | <20 |
| ***21-319-1603*** | 75 | M | 2 | 1,8 | TAF+FTC+BIC | 417 | 0,5 | 0 |
| ***21-319-2348*** | 64 | F | 20 | >4 | TDF+FTC+NVP | 837 | 0,7 | 0 |
| ***21-322-1687*** | 73 | F | 34 | >4 | ETR+RAL | 751 | 1,1 | 0 |
| ***21-322-2289*** | 30 | M | 6 | >4 | DOL+RPV | 869 | 1,6 | 22 |
| ***22-046-2251*** | 62 | M | 32 | >4 | DOL+3TC | 366 | 0,7 | 0 |
| ***22-046-2260*** | 62 | M | 18 | >4 | DOL+3TC | 1070 | 0,7 | 0 |
| ***22-048-1580*** | 31 | F | 7 | >3 | DOL+3TC | 531 | 1,5 | 0 |
| ***22-048-2292*** | 52 | F | 30 | >4 | TAF+FTC+EVG+cobi | 886 | 2,2 | <20 |
| ***22-055-2163*** | 57 | F | 29 | 3 | RPV+CAB-LA | 818 | 2,1 | <20 |
| ***22-059-2522*** | 57 | M | 26 | >4 | NVP+3TC+ABC | 474 | 0,7 | 0 |
| ***22-059-2567*** | 51 | F | 29 | >4 | 3TC+DOL | 419 | 1,2 | 0 |
| ***22-080-1467*** | 57 | F | 32 | >4 | NVP+ABC+3TC | 361 | 1,1 | 0 |
| ***22-080-1521*** | 47 | F | 12 | >4 | DOL+RPV | 587 | 1,3 | 0 |
| ***22-083-1980*** | 48 | M | 17 | 1 | ABC+3TC+RPV/RPV-LA | 458 | 2 | 0 |
| ***22-090-1276*** | 33 | M | 7 | >4 | DOL+RPV | 375 | 1 | 0 |
| ***22-090-2338*** | 76 | M | 18 | >4 | DOL+ABC+3TC | 955 | 1,2 | 0 |
| ***22-132-1260*** | 50 | F | 20 | >5 | DOL+ABC+3TC | 284 | 0,6 | 0 |
| ***22-139-1211*** | 62 | M | 22 | >5 | TAF+FTC+BIC | 709 | 1,2 | 0 |
| ***22-139-1382*** | 76 | M | 28 | >4 | TAF+FTC+BIC | 764 | 1,9 | 28 |
| ***22-327-1600*** | 58 | M | 34 | >5 | DOL+3TC | 409 | 0,6 | 0 |
| ***22-327-2569*** | 22 | F | 6 | >2 | TAF+FTC+EVG+cobi | 622 | 1,1 | <20 |
| ***22-341-2797*** | 51 | M | 12 | >5 | TAF+FTC+EVG+cobi | 924 | 1,2 | 0 |
| ***23-107-5093*** | 67 | M | 22 | >2 | RPV+FTC+TAF | 516 | 0,6 | 0 |
| ***23-108-2607*** | 61 | M | 32 | >5 | DOL+RPV | 525 | 1 | <20 |
| ***23-108-1396*** | 53 | F | 21 | >5 | CAB-LA+RPV-LA | 1044 | 3 | <20 |
| ***23-108-1604*** | 58 | F | 29 | >5 | TAF+FTC+BIC | 1054 | 2,6 | 0 |

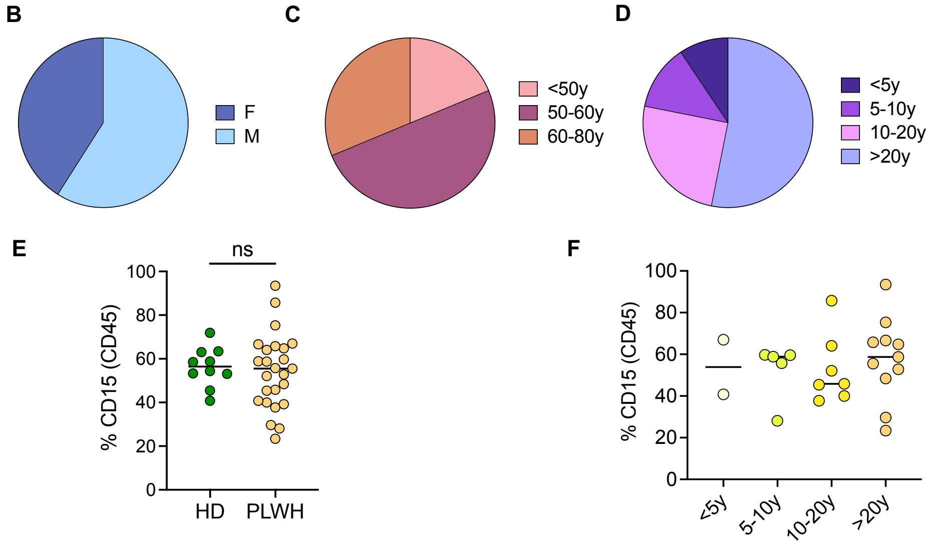


**Supplementary Figure 3. Characteristic of PLWH.**
(A) Demographic and clinical data of ART-treated PLWHIV cohort. (B-C) Distribution in terms of gender (B), age (C) and years since HIV-1 infection diagnosis of PLWH (D). (E-F) Frequencies of neutrophils in blood samples of PLWH (n=25), compared to healthy individuals (n=10) (E) and discriminated by years since diagnosis (F). DOL: Dolutegravir; ABC: abacavir; 3TC: lamivudine; TAF: tenofovir alefenamide; FTC: emtricitabine; BIC: bictegravir; RAL: raltegravir; DRV: darunavir; RTV: ritonavir; EVG: elvitegravir; Cobi: cobicistat; LPV: lopinavir; TDF: tenofovir; NVP: nevirapine; ETR: etravirine; RPV: rilpivirine; CAB: cabotegravir. Suffix -LA means the antiretroviral comes as injections.

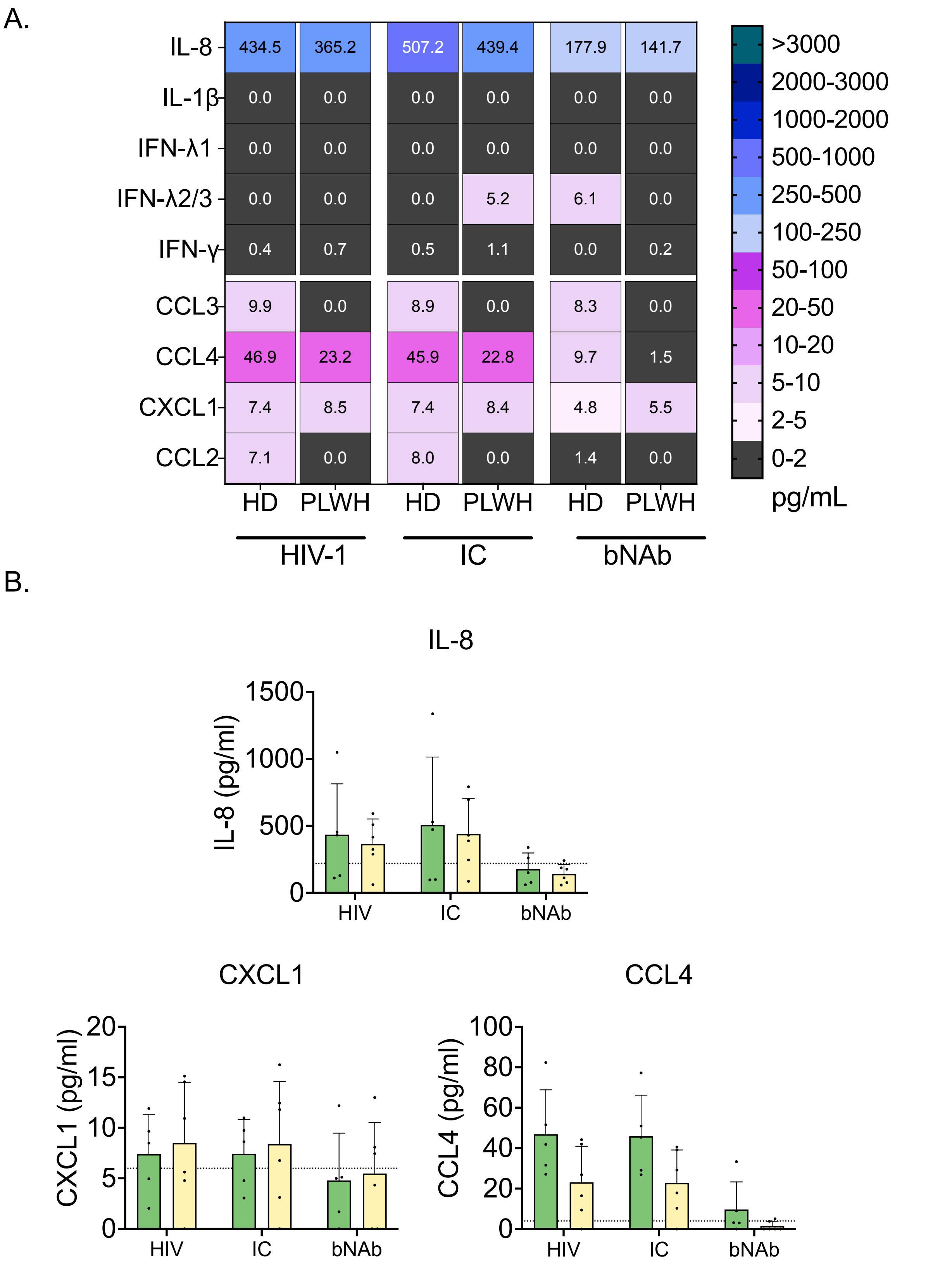


**Supplementary Figure 4. Activation of neutrophils isolated from HD or PLWH stimulated *in vitro* by HIV-1 virions, ICs or free bNAb.** (A-B) Cytokines and chemokines secretion profile, represented as a heatmap (A) and histograms (B) for cytokines/chemokines detected in HD and PLWH. Values are mean +/- SD of 5 independent measurements, including 5 samples of HD and 6 sample from PLWH.

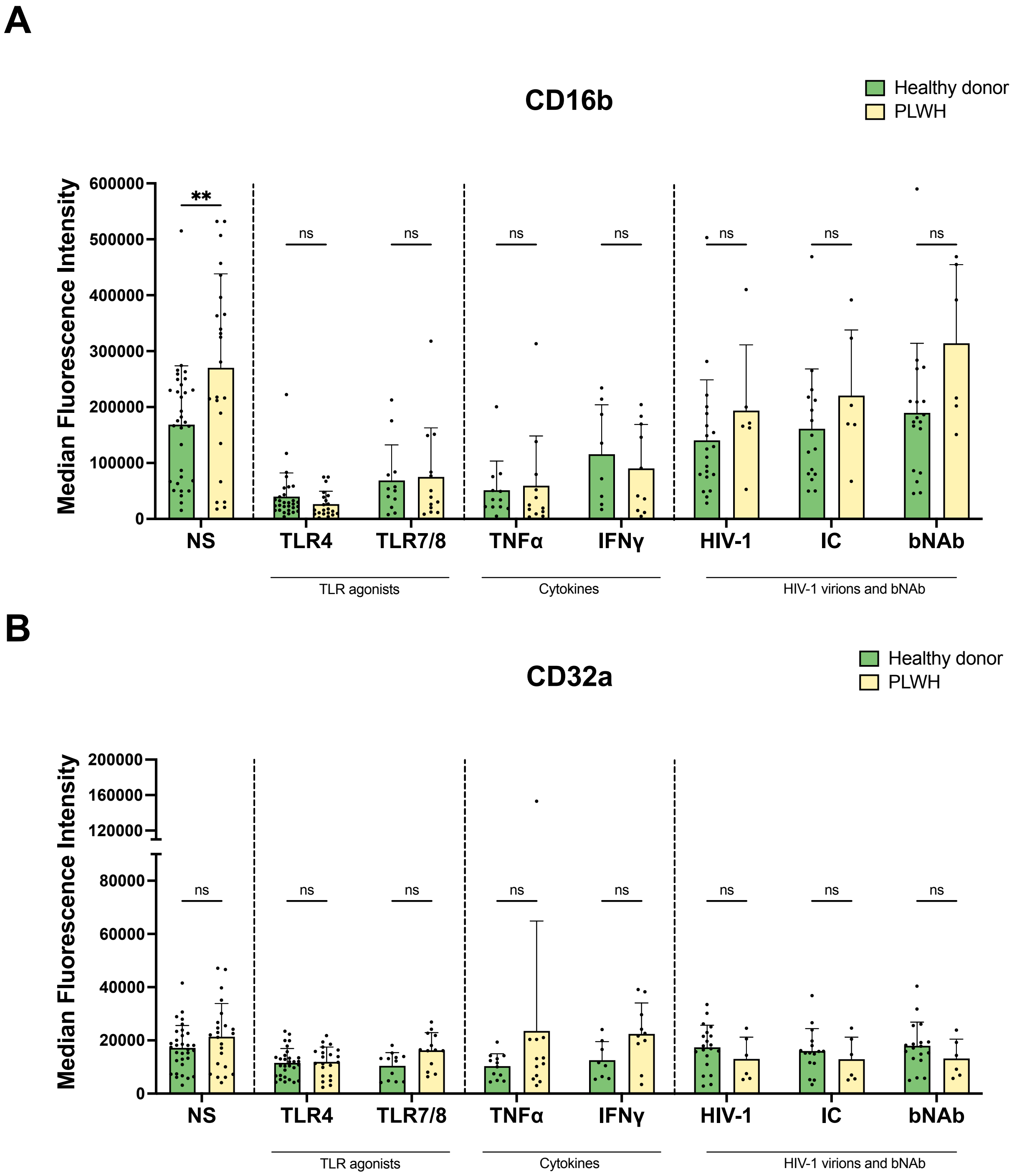


**Supplementary Figure 5. Expression of FcγR on human neutrophils isolated from HD or PLWH at baseline and after stimulations by different stimuli. TLE agonists, pro-inflammatory cytokines, virus.** (A-B) Cell surface levels of CD16 (A) and CD32 (B) upon activation by TLR agonists, cytokines and HIV-1 virions, ICs or free bNAb. Data is represented as MFI Values are mean +/- SD of at least 11 independent measurements. Significance was assigned as follows: ∗ ∗∗p < 0.01

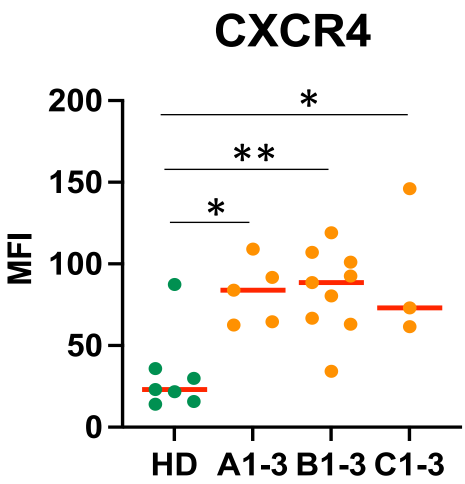


**Supplementary Figure 6. Expresion of CXC4 in neutrophils from HD and PLWH.** CXCR4 expression levels on neutrophils from HD and PLWH (as described in Supplementary Figure 3) are shown at different disease stages, represented as mean fluorescence intensity (MFI). A1-3= 5, B1-3= 9, C1-3= 3.
